# Individual patterns of functional connectivity in neonates as revealed by surface-based Bayesian modeling

**DOI:** 10.1101/2023.07.24.550218

**Authors:** Diego Derman, Damon D. Pham, Amanda F. Mejia, Silvina L. Ferradal

**Affiliations:** Department of Intelligent Systems Engineering, Indiana University, USA; Department of Statistics, Indiana University, USA

**Keywords:** neonatology, brain development, fMRI, RSFC, functional connectivity, resting-state networks, statistical methods, Bayesian modeling

## Abstract

Resting-state functional connectivity is a widely used approach to study the functional brain network organization during early brain development. However, the estimation of functional connectivity networks in individual infants has been rather elusive due to the unique challenges involved with functional magnetic resonance imaging (fMRI) data from young populations. Here, we use fMRI data from the developing Human Connectome Project (dHCP) database to characterize individual variability in a large cohort of term-born infants (N = 289) using a novel data-driven Bayesian framework. To enhance alignment across individuals, the analysis was conducted exclusively on the cortical surface, employing surface-based registration guided by age-matched neonatal atlases. Using 10 minutes of resting-state fMRI data, we successfully estimated subject-level maps for fourteen brain networks/subnetworks along with individual functional parcellation maps that revealed differences between subjects. We also found a significant relationship between age and mean connectivity strength in all brain regions, including previously unreported findings in higher-order networks. These results illustrate the advantages of surface-based methods and Bayesian statistical approaches in uncovering individual variability within very young populations.

## 1 Introduction

The human brain undergoes significant changes during late gestation and early infancy. Characterizing the functional brain organization during early postnatal ages is clinically relevant to elucidate how different genetic factors and environmental hazards may impact subsequent development. Resting-state functional connectivity (RSFC) is a widely used approach to delineate the functional brain organization in the perinatal period since subjects are imaged at rest without requiring the performance of any specific task. Typically, RSFC methods look for temporal coherence of spontaneous fluctuations of functional magnetic resonance imaging (fMRI) signals to define resting-state networks (RSNs), also known as the functional connectome. Seminal studies based on RSFC methods have shown that the functional connectome during early development is affected by premature birth (Doria et al., 2010; Smyser et al., 2010), maternal stress (DeSocio, 2018; Sandman et al., 2012), developmental dyslexia (Cao et al., 2014; Yu et al., 2022), and prenatal drug exposure (Merhar et al., 2021), among other factors. While these studies have provided important insights into the typical and atypical functional brain organization in early life, the vast majority have focused on the estimation of RSNs at the group level (i.e., derived from multi-subject data). In recent years, it has been shown that the functional connectome is associated with later cognitive and behavioral outcomes (Ali et al., 2022; He et al., 2018; Keller et al., 2023; Newbold et al., 2020; Rosenberg and Finn, 2022). Therefore, individual characterizations are critical to making subject-based predictions of clinical relevance.

The robust estimation of RSNs in individual infants has been rather elusive due to the inherently low signal-to-noise ratio (SNR) of the resting-state fMRI data and the unique sources of noise (e.g. idiosyncratic head motion, faster cardiac and respiration rates) in this population (Spann et al., 2023). Traditional RSFC analysis approaches like group independent component analysis (gICA) produce clean estimations of RSNs when combining data from a large number of infants (Eyre et al., 2021; Gao et al., 2015) but fail to capture individual differences. In contrast, estimations of subject-level RSNs obtained from methods such as dual regression have low power to mitigate noise from single-subject data. Precision functional mapping is an emerging technique that offers an effective strategy to increase the SNR in functional connectivity analysis within individuals, maximizing the power to detect individual differences in RSNs (Gordon et al., 2017; Gratton et al., 2022, 2020; Laumann et al., 2015). However, to achieve an acceptable level of reliability at the subject level, extensive amounts of imaging data need to be collected, which is rarely practical or cost-effective when considering infant and clinical populations. Bayesian approaches offer a powerful complement to longer scans by leveraging shared information across subjects from a representative population, which reduces noise while enabling individual differences to be expressed without requiring extended scans (Kong et al., 2019; Mejia et al., 2020).

In this study, we explore hierarchical Bayesian modeling in combination with surface-based analysis to improve individual estimations of RSNs in infants during the first weeks of postnatal life. The proposed Bayesian framework (Mejia et al., 2020) requires population-derived priors or templates, so we leverage the information contained in the large open-access dataset from the developing Human Connectome Project (dHCP) (Edwards et al., 2022). A critical assumption of the Bayesian framework is that all the subjects are anatomically co-registered to a common atlas space. Thus, to optimize the anatomical alignment across individuals with different gestational ages, the analysis is entirely done on the cortical surface using registration techniques guided by age-matched infant atlases. While surface-based registration approaches have become increasingly popular in studies of functional connectivity involving adult populations (Glasser et al., 2016), their application to early brain development is just now gaining traction. Notably, Wang et al., 2023 were able to generate cortical parcellation maps in a cohort of infants/toddlers by leveraging local FC gradient maps that improve functional alignment across subjects after folding-driven registration. In addition, Hu et al., 2022 used the same folding-driven registration to demonstrate the existence of functional networks in neonates, and their longitudinal stability throughout the first two years of life.

Our analysis of a cohort of 289 term-born neonates (age at scan: 37.4 - 44.8 weeks) shows that the surface-based hierarchical Bayesian approach can produce clean estimates of subject-level cortical RSNs using only 10 minutes of individual fMRI data. The statistical framework enables the computation of subject-level t-statistic maps which, in turn, allows for functional parcellations at the subject level. As a result, we observed individual topographical differences hidden by group-level averages. Furthermore, a positive relationship between age and subject-level connectivity strength was revealed for almost all cortical RSNs, confirming the hypothesis that the functional connectome matures with age in infants. Importantly, our work extends beyond existing research by showing that significant maturational changes are not only restricted to primary sensory networks but also present in higher-order networks in the first weeks of postnatal life. These results illustrate the advantages of combining surface-based processing and hierarchical Bayesian approaches to inform individual variability in very young populations, opening the door to precision neuroimaging studies of early brain development with enhanced accuracy and reliability.

## 2 Methods

### 2.1 Subjects and data acquisition

MRI data was obtained from the second release of the dHCP database. For this study, we only considered term-born infants (i.e. gestational age (GA) >= 37 weeks) with radiological scores lesser than three, indicating no lesions of clinical or analytical significance (see dHCP release notes^1^ for details). Following these criteria, 305 term-born infants (age at birth: 37.1 – 42.3 weeks gestational age, GA) scanned shortly after birth (age at scan: 37.4 – 44.8 weeks postmenstrual age, PMA) were considered for further analysis.

All scans were obtained with a 3T Philips Achieva using a dedicated neonatal head coil at Evelina Newborn Imaging Centre, St. Thomas Hospital, London, UK. Both T1-weighted (TR = 4795 ms; TE = 8.7 ms) and T2-weighted (TR = 12 s; TE = 156 ms) structural scans were obtained with a multi-slice Turbo Spin Echo (TSE) sequence, with in-plane resolution 0.8 x 0.8 *mm*² and 1.6 *mm* slices overlapped by 0.8 *mm*. Two stacks of images were taken per weighting, sagittal and axial, which were integrated to obtain T2w volumes with an image resolution of 0.8 mm isotropic. Blood oxygen level-dependent (BOLD) scans were obtained with a multi-slice gradient-echo echo planar imaging (EPI) sequence (TE = 38 ms; TR = 392 ms, multiband factor = 9; flip angle = 34°) with an image resolution of 2.15 mm isotropic. A resting-state BOLD fMRI acquisition of 2300 time points (15 minutes) was obtained for each infant.

### 2.2 MRI preprocessing

#### 2.2.1 Cortical surfaces

After brain extraction using a modified version of FSL BET for unmyelinated brains, tissue-specific masks corresponding to gray matter, low-, and high-intensity white matter were segmented from the brain-extracted T2w images using the Draw-EM tool (Makropoulos et al., 2014). Individual surfaces corresponding to white matter, pial, and midthickness were created using these tissue masks as described in Makropoulos et al., 2018.

For the rest of the analysis, a 40-week symmetrical atlas consisting of 57,700 vertices after excluding the medial wall built from the dHCP cohort (Williams et al., 2023) was used. The original atlas in Bozek et al., 2018 was built based on a spherical registration approach using the Multimodal Surface Matching (MSM) tool for aligning cortical folding (Robinson et al., 2018). For this extended version, left-right vertex correspondence was enforced by co-registering right and left sulcal depth maps during alignment.

We used the MSMSulc approach to register each individual cortical surface to the symmetrical 40-week atlas, concatenating single-week transformations to avoid the accumulation of approximation errors. MSMSulc uses a highly regularized folding-based registration that has been shown to improve functional overlap among subjects (Robinson et al., 2018). These transforms were calculated using the scripts from the dHCP GitHub repository ^2^. The spherical registrations from this process were later used when mapping the BOLD fMRI data onto the atlas surface.

#### 2.2.2 Functional data

Resting-state BOLD fMRI data were preprocessed using a modified version of the Human Connectome Project (HCP) (Glasser et al., 2013) pipeline specifically developed for the dHCP (Fitzgibbon et al., 2020).

Preprocessing steps were performed on the BOLD volumes in native space and included: (1) field map correction, (2) intra- and inter-volume motion correction, (3) high-pass filtering to remove slow drifts, and (4) spatial ICA using FSL FIX (Griffanti et al., 2014; Salimi-Khorshidi et al., 2014) to regress out structured noise artifacts.

To further reduce the potential effect of motion on functional connectivity measures, we adopted a conservative frame censoring approach. Emerging evidence suggests that head motion is correlated with neurophysiological states (e.g., sleep state) (Denisova, 2019). Since aggressive frame censoring may introduce heterogeneities in the BOLD time series associated with different physiological states, we retained a contiguous block of data to minimize this potential issue. Following the criteria proposed in Eyre et al., 2021, frames with DVARS (root-mean-squared BOLD signal intensity change) higher than 1.5 times the interquartile range above the 75th percentile within a session were considered corrupted by motion. A contiguous block of 1600 frames (∼10 minutes) with the minimum number of motion outliers was retained for each subject. Furthermore, any subject with more than 160 motion-corrupted frames (10%) within the contiguous block was excluded. By following this approach, we were able to retain 274 subjects with 10 minutes of resting-state BOLD fMRI data for further analysis.

The volumetric resting-state BOLD data was mapped onto the individual cortical surfaces using the HCP surface pipeline (Glasser et al., 2013) as implemented in Connectome Workbench. The surface-based BOLD data was then mapped onto the custom 40-week atlas surface using spherical registration (as described in §2.2.1 *Cortical surfaces*). Finally, the individual datasets were spatially smoothed using a geodesic 2D Gaussian kernel (FWHM = 3 mm). In the following analyses, only the cortical data was taken into consideration.

#### 2.2.3 Temporal signal-to-noise ratio

To quantify image quality throughout the cortex and mask out noisy vertices, we computed a measure of temporal signal-to-noise ratio (tSNR) as the ratio between the mean BOLD signal and its standard deviation. A global tSNR map was built in decibels [*dB*] by averaging the individual tSNR maps across the whole sample as

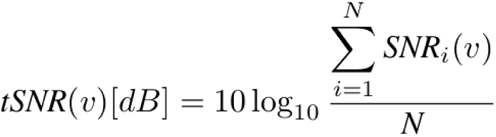

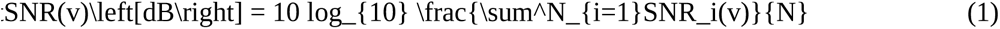

where, for each subject *i*, the SNR is calculated for each vertex *v*.

As expected from the patterns associated with susceptibility artifacts in BOLD data and previous similar analyses (Fitzgibbon et al., 2020; Yeo et al., 2011), we observe areas of particularly low tSNR in the inferior and medial temporal lobe, orbitofrontal and insular cortices (Fig. S3).

### 2.3 Bayesian subject-level ICA

To estimate subject-level RSNs, we use a custom implementation of template ICA (Mejia et al., 2020), an extension of probabilistic ICA (Beckmann and Smith, 2004). In this section, we describe the template-based ICA model, the calculation of population-derived priors, and the strategy for estimating individual posterior mean and variance.

#### 2.3.1 Template-based ICA model

For each subject *i*, the dimensionally reduced BOLD fMRI data **y***_i_* with *T* time-points is modeled at vertex *v* as

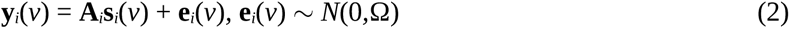

where **A**^(*T×Q*)^ is the mixing matrix, **s***_i_*^(*Q*×*1*)^ represents the *Q* independent components (ICs) associated with a different RSN, and **e***_i_*^(*T*×*1)*^ represents any source of Gaussian noise with variance Ω. Like in probabilistic ICA, **y***_i_*is obtained using single value decomposition (SVD) of the preprocessed BOLD dataset, and an estimate of the Gaussian noise variance **Ω** is obtained as the residual variance. Note that the error in equation 2 has a Gaussian distribution since we assume that any source of non-Gaussian noise is eliminated after ICA-FIX on each preprocessed BOLD dataset. In the template ICA model, we further characterize each IC as

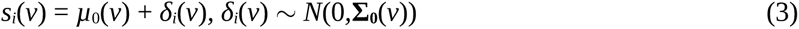

where *µ*_0_(*v*)^(*Q*×1)^ is the group-level mean, and *δ_i_*(*v*)^(*Q*×1)^ represents subject-level deviations denoting unique features of spatial topography associated with each RSN. We assume that the *δ_i_*(*v*) are normally distributed with covariance matrix 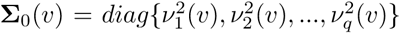 where each 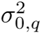 is the between-subject variance associated to each IC *q* at each vertex *v*.

For a subject-specific BOLD dataset *y_i_*(*v*), the proposed problem can be formulated within a hierarchical Bayesian framework where we aim to obtain a vertex-wise posterior distribution for the subject-level IC maps *s_i_*(*v*), given a set of population priors or “templates” (mean *µ*_0_(*v*) and between-subject variance **Σ**_0_(*v*)). Note that given the Gaussian prior and likelihood, the vertex-wise posterior spatial maps are Normally distributed with a subject-level mean *µ_i_* and variance *σ_i_* that have analytical forms. To estimate the posterior mean and variance of the subject-level ICs, along with the model parameters including the mixing matrix and residual variance, a computationally efficient expectation-maximization (EM) algorithm is utilized (see §2.3.3 *Estimation of individual posterior mean and variance*) and Figure 1 for a description of the estimation procedure).

**Figure 1:**
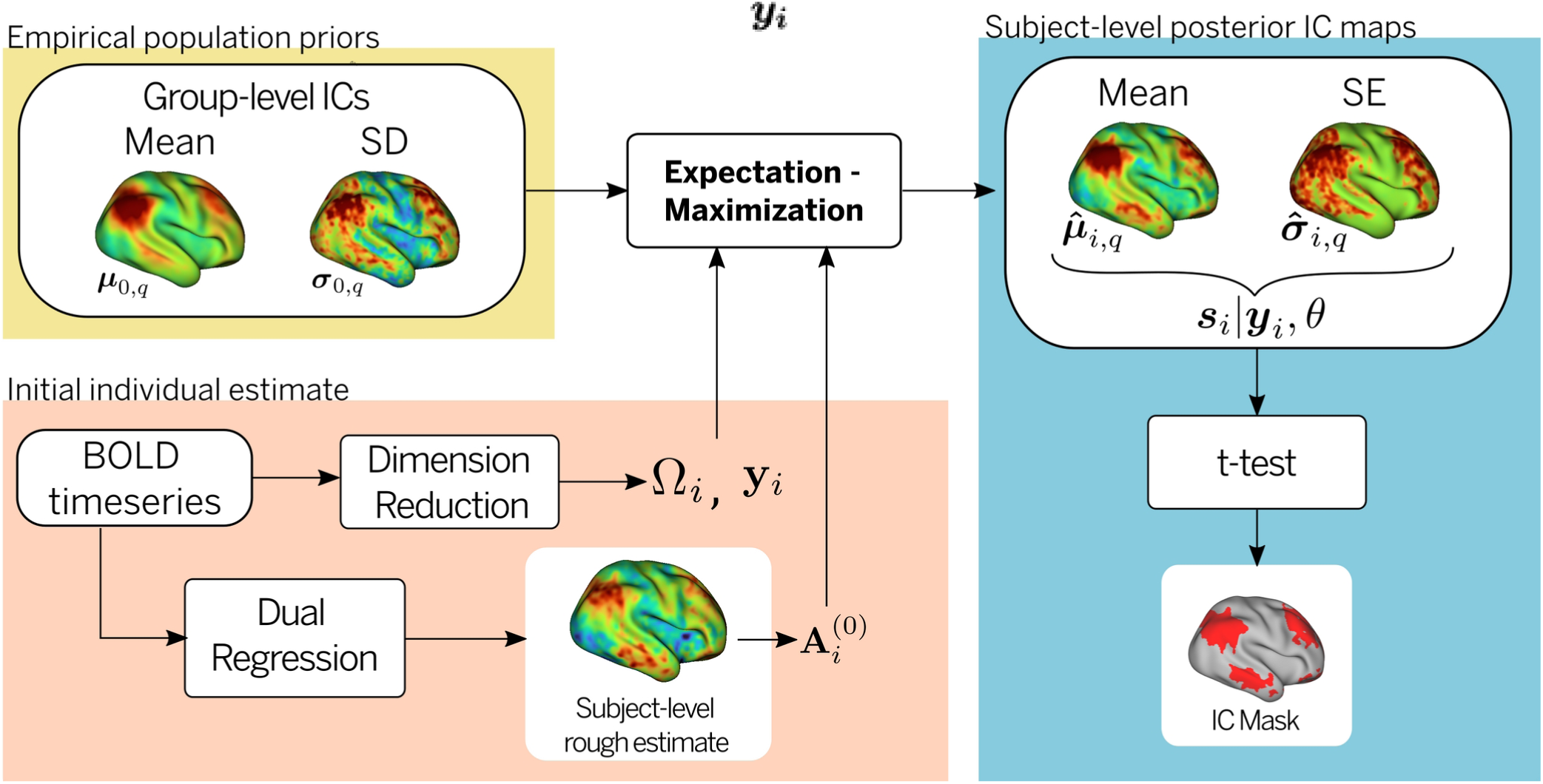
Estimation of subject-level IC maps. To estimate the posterior mean and variance of the subject-level ICs, along with the model parameters including the mixing matrix and residual variance, a computationally efficient expectation-maximization (EM) algorithm is used. An initial estimation of the model parameters (*Ω_i_*) is obtained from dimensionality reduction ( *y_i_*) of the BOLD timeseries, while the mixing matrix 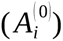 is obtained from a rough dual regression estimate of the individual IC maps. The empirical population prior parameters, namely, the mean ( *μ*_0_ (*v* )) and the between-subject variance *σ* _0_( *v* ) of the group-level IC maps are calculated from a uniform sample of the cohort. After the EM algorithm converges, an estimate of the subject-level mean 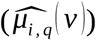 and standard deviation 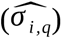 for each IC is obtained. A binary mask of significant engagement for each IC is obtained from a t-test corrected for multiple comparisons on the individual mean and variance IC maps.

#### 2.3.2 Population priors

The Bayesian framework relies on population-derived priors, which consist of estimates of the mean and between-subject variance of a set of ICs. To first define the set of ICs, we perform group ICA (Beckmann et al., 2005) on a subset of 24 subjects (scanned between 43.5 and 45 weeks PMA) using FSL MELODIC (C. Beckmann and Smith, 2004). The number of ICs is set at 20, excluding subcortical regions, to achieve a balance between robustness and similarity to previous analysis on infants (Doria et al., 2010; Eyre et al., 2021; Rajasilta et al., 2020; Toulmin et al., 2015). Eight ICs were associated to neurologically relevant cortical networks, namely, medial motor, lateral motor, auditory, somatosensory, primary visual, motor association, visual association, and default mode network (DMN) (Fig. 4). Examples of the timeseries and power spectrum associated with signal and nuisance components is available in Fig. S6. To obtain noisy pseudo test-retest point estimates of individual IC maps that can be used to obtain population means and between-subject variance estimates, dual regression (Nickerson et al., 2017) is applied to a subset of subjects (N = 35) uniformly scanned between 37 and 43 weeks PMA. To calculate the between-subject variability, each individual BOLD time series is divided into two halves (or pseudo-sessions) prior to dual regression. The population mean *µ*_0_(*v*) for each IC *q* is estimated as

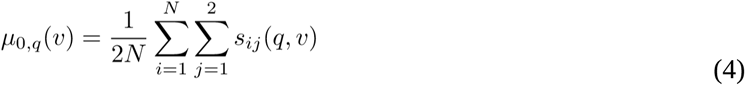

where *s_ij_*(*q,v*) is the dual regression IC estimate for each subject *i* and each session *j*.

The between-subject variance is obtained by decomposing the total variance into within- and between-subject components. The total variance for each individual IC at each vertex was estimated as the average of the variance obtained for each pseudo-session,

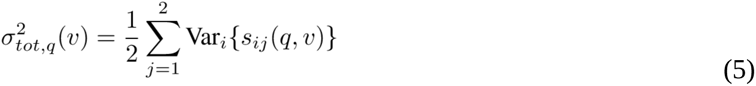

while the within-subject variance can be estimated based on the variance of the difference between the individual ICs from each pseudo-session

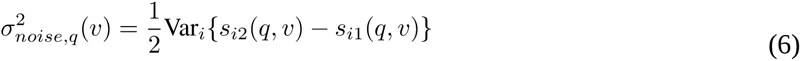

Finally, from equations 5 and 6, the between-subject variance is estimated as the difference between the total and within-subject variances

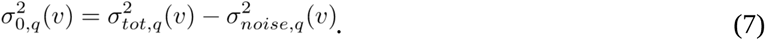

where the *σ*_0*q*_ are the diagonal elements of the covariance matrix **Σ**_0_ in equation 2, representing the variance of the empirical population prior.

#### 2.3.3 Estimation of individual posterior mean and variance

A schematic of the estimation procedure is shown in Figure 1. The EM algorithm requires an initial estimate of the parameters **Ω** and **A_i_**. **A_i_^(0)^** was obtained from the dual regression procedure described above. Gaussian noise variance 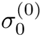 was estimated from the residual variance of the dimensionality reduction of the BOLD data.

These two parameters are updated at every iteration of the EM algorithm by maximizing the expected log-likelihood. Alternately, conditional on the current estimate of the parameters, the posterior distribution of the subject-level ICs **s_i_** is updated to obtain the posterior moments required for the parameter maximum likelihood estimations. The algorithm runs until convergence of the parameter estimators, and the posterior mean *µ***^***_i_* and variance *σ***^***i* of *s*^*i*(*v,y,θ*) are obtained. A schematic representation of the entire pipeline is shown in Figure S1.

## 3 Results

### 3.1 Individual patterns of functional connectivity

Using only 10 minutes of resting-state fMRI data, we identified eight cortical RSNs (represented by independent components, ICs) in individual subjects from a cohort of 239 full-term infants (note that the 35 subjects considered in the estimation of the empirical population priors were excluded from further analysis). Overall, the surface-based Bayesian approach produced cleaner estimates of subject-level IC maps than dual regression, as illustrated by five representative IC maps obtained in a single infant (Fig. 2). Notably, the Bayesian approach removed clusters of spurious activations present in the dual regression maps (see for example the temporal cortex in the somatosensory IC maps of Figure 2). Other improvements include the emergence of more defined clusters in distributed networks, as shown in the prefrontal cluster of the default mode IC maps. A complete set of subject-level IC maps obtained for both methods is included in Figure S2.

**Figure 2:**
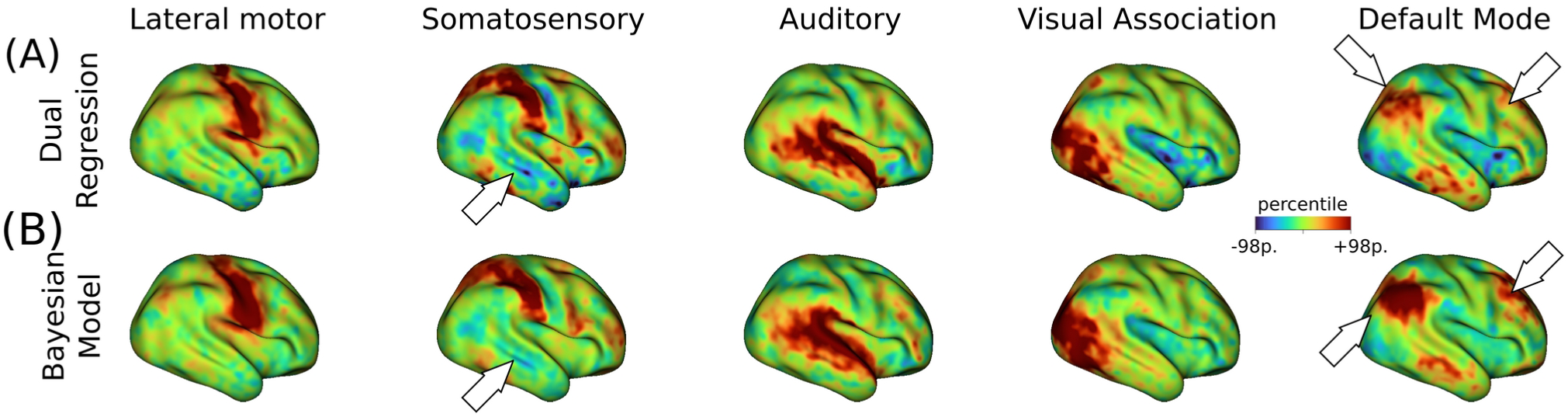
Subject-level maps obtained from 10 minutes of resting-state fMRI data. (A) Five representative IC maps obtained from dual regression for a term-born infant scanned at 42.6 weeks PMA. (B) Posterior mean IC maps derived from the Bayesian model produced cleaner maps than the dual regression approach. White arrows highlight some areas of notable improvement in cluster convexity (e.g. default mode network) or reduction of spurious engagement (e.g. somatosensory, inferior parietal). All maps are projected onto the inflated 40-week atlas. A symmetrical scale around zero is defined by setting the maximum at the 98th percentile of FC (a.u.) for each IC map.

To perform comparisons across subjects, we computed t-statistic maps from the subject-specific posterior mean and standard error maps estimated by the surface-based Bayesian model. Figure 3 shows the unthresholded t-statistic maps for five networks and three term-born infants scanned at different postmenstrual ages within a period of four weeks. While a general spatial topography is preserved for each specific network, individual differences are evident across subjects. For example, the temporal cluster of the default mode IC map of subject A is more defined than in subjects B and C. Similarly, the prefrontal cluster of the same IC map of subject B shows a stronger level of engagement in comparison with subjects A and C.

**Figure 3:**
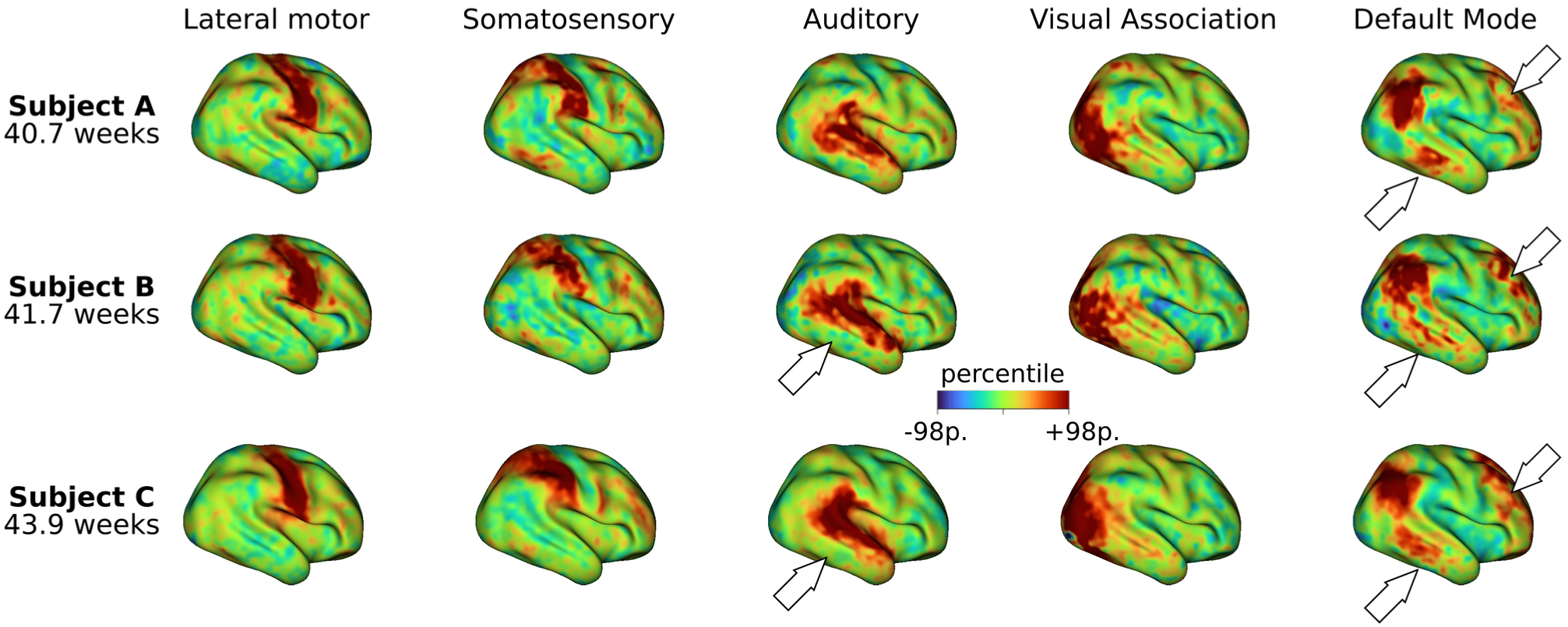
Subject-level maps across different subjects. Unthresholded t-statistic maps were computed from the mean and standard error maps derived from the Bayesian model estimation. The five ICs shown in Figure 2 are displayed for three term-born infants scanned at 40.6 weeks PMA, 41.7 weeks PMA, and 43.9 weeks PMA, respectively. Although a general spatial topology is preserved within each specific network, there are evident variations across individuals (as indicated by the white arrows).

**Figure 4:**
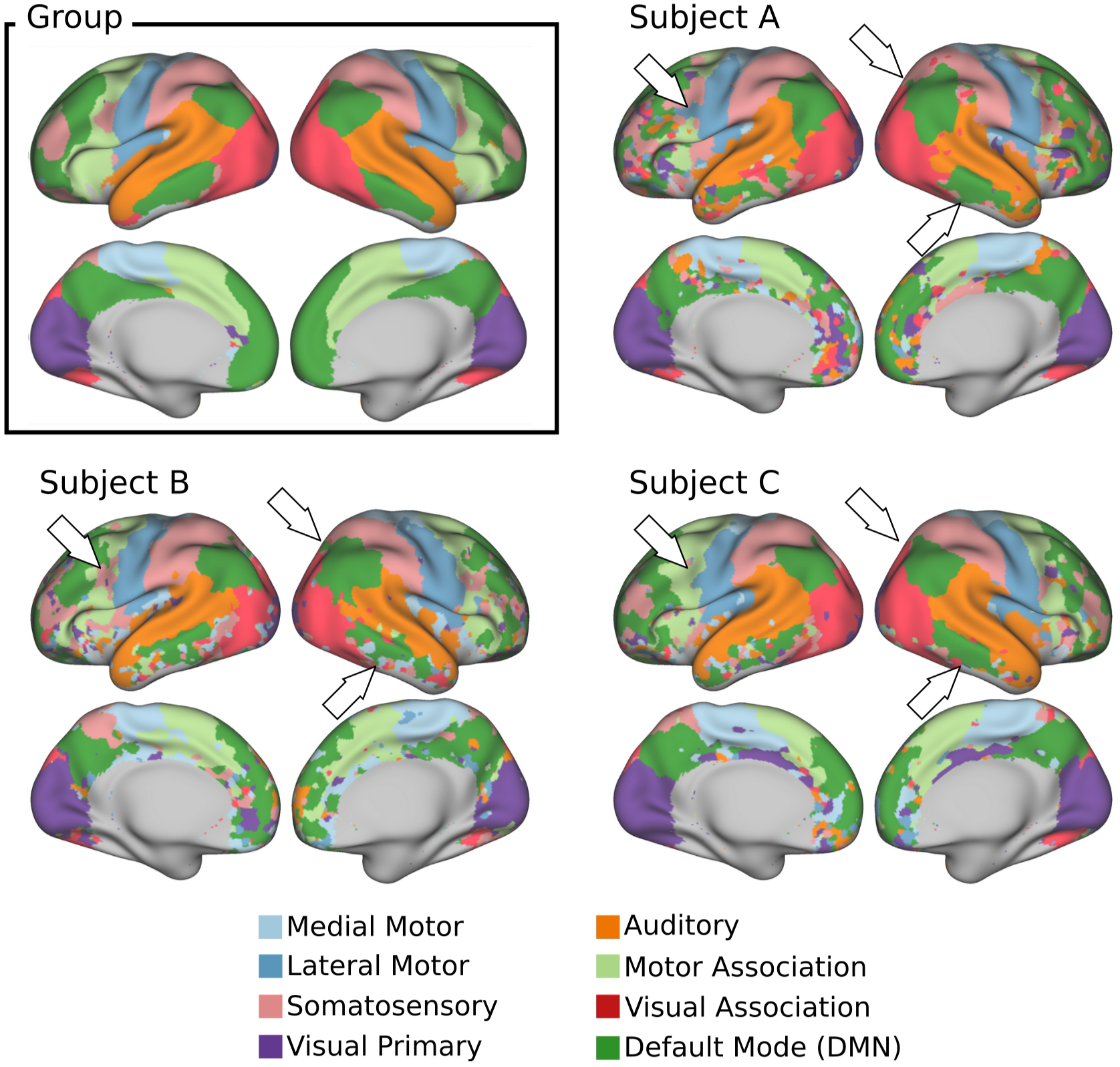
Group vs. individual parcellations. Functional parcellations were obtained using a winner-takes-all approach, dividing the cortex into eight distinct brain networks and subnetworks. The group parcellation was obtained from the Z-score maps derived from the group ICA analysis on 24 subjects. The individual parcellations were obtained from the subject-level t-maps derived from the Bayesian estimation on the subjects shown in Figure 3. White arrows highlight notable topographical differences between subjects. The results are displayed only in regions where the global signal-to-noise ratio (SNR) is greater than 17 dB (see Figure S3 for a whole depiction of the global SNR computed for this cohort). All parecellation are projected onto the 40-week inflated atlas.

As an alternative way to summarize the individual differences observed in Figure 3, we used a winner-takes-all (WTA) strategy to obtain subject-level functional parcellations (Fig. 4) in which a network label was assigned to each vertex based on the highest t-score at that location (see Fig. S5 for a comparison between the individual IC maps and their associated WTA parcellations). The Bayesian approach constitutes a compromise between robust group-level RSNs provided by the population template and subject-level RSNs associated with the individual data. This can be appreciated by looking at the functional parcellation maps obtained for the group ICA maps and the subject-level parcellations for three different infants (Fig. 4). Even though a general resemblance to the group-level parcellation is preserved, remarkable individual differences in shape, size, and location of the individual parcels are evident across subjects. See, for example, the significantly reduced temporal cluster of the default mode IC map in the right hemisphere of Subject B in comparison with the other individuals.

### 3.2 Effect of age at scan

To explore whether the observed differences are related to maturational factors, we computed the Spearman’s rank correlation coefficient between age at scan and individual connectivity strength for each network, after controlling for sex and motion. Statistical significance was defined as monotonic trends with a p-value lower than 0.05, as calculated using the AS 89 algorithm (Best and Roberts, 1975) implemented in the ‘stats’ R-package. Individual connectivity strength was defined as the mean t-score within the areas of significant engagement or IC mask, for each subject-level ICs. Areas of significant engagement were estimated from a t-test on each pair of posterior mean and standard error IC maps after Bonferroni correction for multiple comparisons (as shown in the pipelines of Fig. 1 and Fig. S1). To account for variable vertex area across the cortical surface, the individual connectivity strength was also weighed by the midthickness vertex area within each IC mask. Within-network connectivity significantly increased with age at scan (37.4 – 44.8 weeks PMA) for all cortical ICs, including medial motor, lateral motor, somatosensory, auditory, visual, motor association, default mode, and visual association networks (Fig. 5).

**Figure 5:**
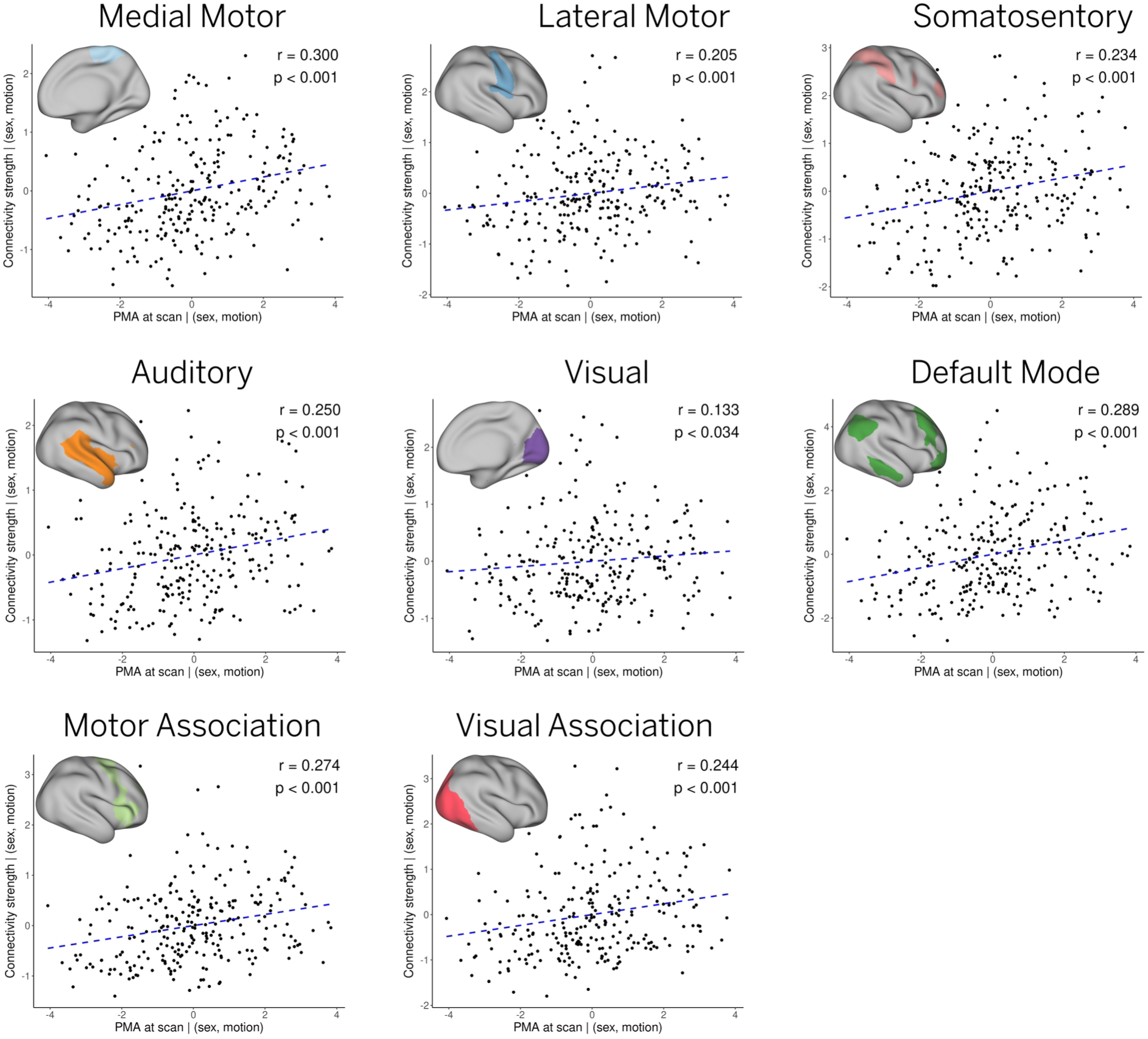
Effect of age at scan. The relationship between age at scan and individual connectivity strength for each network was assessed by a Spearman’s rank correlation test, after controlling for sex and motion. All networks exhibited significant effects with age (p < 0.05) as determined by the corresponding Spearman’s p-values. A dashed blue line showing a linear trend is also included for illustration purposes.

### 3.3 Inter-subject variability

To quantify the spatial variability across individuals, we estimated a frequency map from the subject-level parcellations. Figure 6 shows the percentage of subjects that share the same label or parcel at each cortical vertex. Given that multiple labels may contribute to the same vertex in different subjects, the frequency map only shows the percentage of subjects associated with the dominant label at each location. In addition to all primary networks, the visual association networks and the posterior nodes of the default mode network appear clearly defined in the frequency map and exhibit a high spatial overlap. In contrast, the clusters associated with the prefrontal node of the default mode network and motor association networks exhibit lower spatial overlap across individuals.

**Figure 6:**
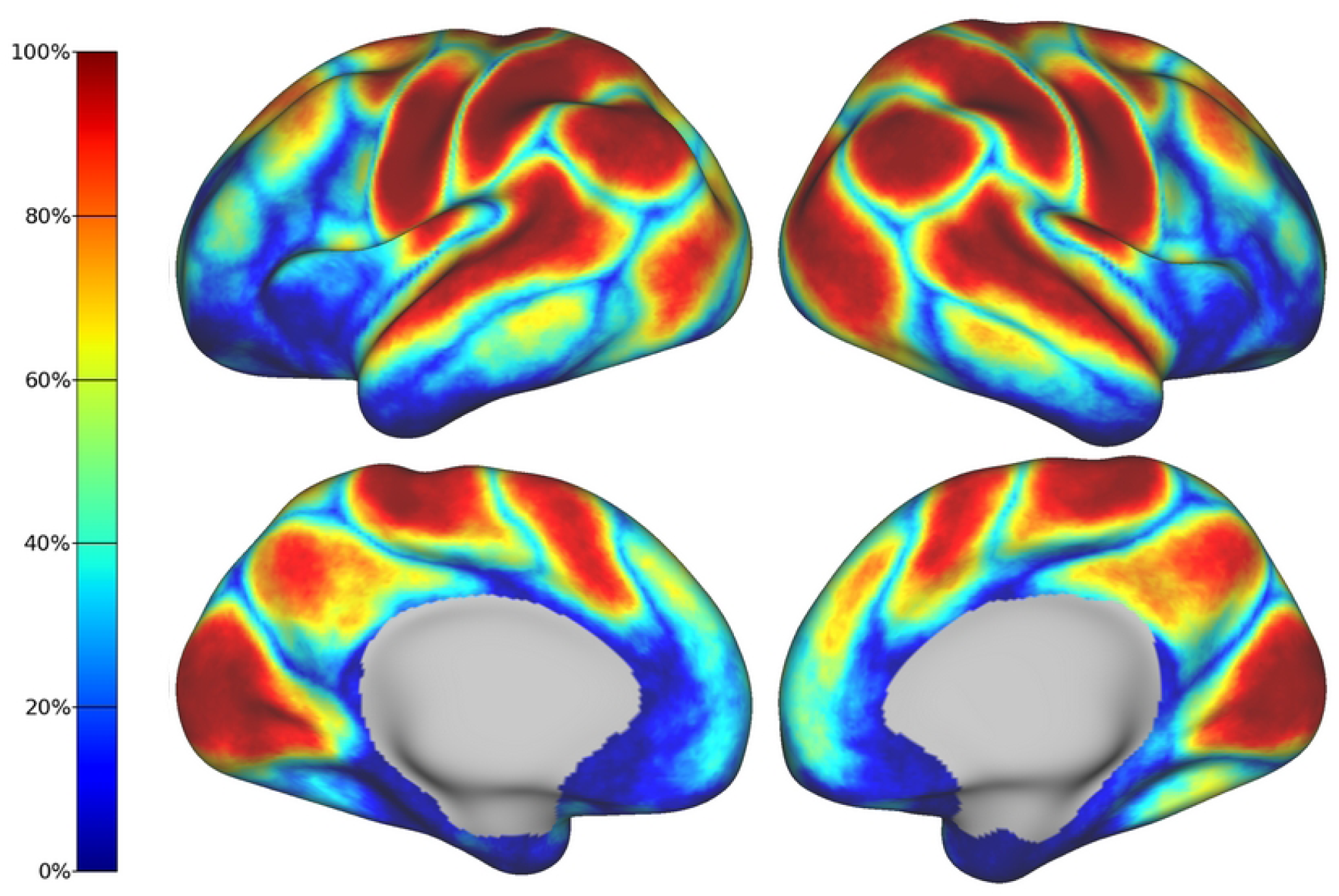
Frequency map of the entire cohort. Each vertex represents the percentage of subjects that share the same parcel or network. Networks with higher spatial overlap such as the primary sensory RSNs are represented by warm colors. Note that boundaries between networks and areas with low SNR show higher variability (i.e., low spatial overlap) across subjects.

## 4 Discussion

This study extends beyond existing research by characterizing RSNs in term-born neonates at the subject level, leveraging several technical improvements to elucidate individual differences and developmental trajectories within the first weeks of life. Although the existence of resting-state networks (RSNs) is well documented in adults, several neonatal studies have demonstrated the presence of recognizable, albeit sometimes incomplete, RSNs at the group level (Doria et al., 2010; Eyre et al., 2021; Gao et al., 2015; Smyser et al., 2010). Our study presents a more complete picture at the individual level of the functional connectome at the time of birth. To achieve these results, we utilized surface-based methods and age-matched atlases to mitigate co-registration errors and partial volume effects. In addition, we adopted a Bayesian approach that offers a powerful statistical framework to obtain clean estimates of subject-level RSNs using a limited amount of individual data but borrowing strength from population-derived priors.

Using these methodological innovations, we were able to identify topographical differences across subjects hidden by group averages. This is in line with adult studies showing that areas of common activation obscure individual connectivity features (Gordon et al., 2017; Hermosillo et al., 2022; Michon et al., 2022). This is especially important because individual parcellations derived from individual connectivity estimations are susceptible to suffering from coregistration misalignment between subjects (Bijsterbosch et al., 2019). Indeed, inter-subject differences are evident across the individual t-maps (Fig. 3) and functional parcellations (Fig. 4) obtained for different infants. Interestingly, the individual parcellations also show topographical differences with respect to the group-level parcellation. As an example, note the individual differences in the temporal and prefrontal clusters of the default mode network; while the group presents a single representation (Fig. 4 Group), different subjects present variable cluster shape and sizes (Fig. 4 A-C). It is notable that despite these differences, the results show adult-like higher-order RSNs maps already present at birth – albeit sometimes in sparser, precursory form (Fig. 3).

To quantitatively assess these individual differences, we studied the relationship between subject-level RSNs and age at scan. As in previous reports, significant maturation is observed in all primary sensory RSNs, as measured by the correlation between network strength and age at scan (Fig. 5) (Doria et al., 2010; Eyre et al., 2021; Gao et al., 2015; Hu et al., 2022; Smyser et al., 2010). Furthermore, contrary to prior studies, we also note a clear developmental trajectory within a five-week window for the independent components (ICs) linked to higher-order RSNs.

The frequency maps depicting the spatial overlap of networks across subjects reveal a pattern that progresses from posterior to anterior regions (Fig. 6). In particular, we observe high levels of spatial overlap in the primary sensory networks, along with the higher-order networks located in the posterior regions. In contrast, the prefrontal cluster of the DMN displays low spatial overlap which can be attributed to developmental factors, as supported by histological findings (Keunen et al., 2017). Notably, there are other regions characterized by low spatial overlap, namely, the boundaries between the different RSNs as well as the insular and inferior temporal cortices. The lack of consistency in the delineation of boundaries between the RSNs reflects the topographical variability that our study aims to capture. By contrast, the limited overlap observed in the insula and inferior temporal cortices may be associated with elevated levels of noise in the BOLD signal, as demonstrated by the tSNR map obtained for this specific cohort (Fig. S3).

### 4.1 Subject-level inference produces more accurate descriptions of brain organization

We hypothesize that variability in the spatial distribution of RSNs across individuals could be the primary factor that hinders the identification of population-level trends and renders growth curves challenging to obtain, especially for distributed higher-order RSNs. In contrast to prior studies, the use of individual regions of interest (ROIs) enabled us to achieve improved outcomes in terms of estimating within-network connectivity strength at the individual level. This is consistent with previous reports showing group-average ROIs that lead to inflated or under-estimated group differences (Levi et al., 2023).

The trend towards individual inference instead of group analysis has emerged as a recent focus in precision neuroimaging. This approach facilitates the acquisition of relevant individual insights in the clinic and also plays a critical role in aggregating individual data without diluting individual features (Gordon et al., 2017; Gratton et al., 2020; Laumann et al., 2015). To achieve acceptable levels of reliability at the individual level using naive statistical methods, extensive amounts of imaging data need to be collected, which is rarely practical in infants who are typically scanned during natural sleep. As an alternative, we adopt a recently proposed empirical Bayesian model (Mejia et al., 2020) that leverages the growing volume of neonatal data available in existing databases such as the dHCP database. The adopted model produces more accurate subject-level RSN maps by shrinking to the empirical population prior in subject-specific areas of low SNR while maintaining the individual differences.

### 4.2 Advanced techniques are crucial to maintain precision mapping

Concurrently, we used additional technical enhancements to mitigate any biases that could potentially skew our results toward adult populations. To eliminate any distortions that may arise from mapping neonatal brains onto an adult-based atlas, we utilized a symmetrical atlas (Bozek et al., 2018; Williams et al., 2023) derived from the dHCP cohort. This atlas served as a reference surface onto which the individual BOLD timeseries were projected.

Moreover, the adoption of surface-based analysis effectively reduces the potential influence of partial volume effects encountered in volumetric analysis (Coalson et al., 2018). By applying spatial smoothing on cortical surfaces, contamination between functional regions on opposite sides of a sulcus is mitigated (Brodoehl et al., 2020) because the filtering kernel is applied over 2D geodesic distances, which are more neuroanatomically relevant, instead of 3D volumetric distances (Glasser et al., 2013).

Additionally, alignment of the cerebral cortex to the atlas by surface registration algorithms simplifies the 3D problem of volumetric registration to 2D, resulting in improved and more robust co-registration (Robinson et al., 2014). This is especially important in this work because individual parcellations derived from individual connectivity estimations are susceptible to suffering from co-registration misalignment between subjects (Bijsterbosch et al., 2019). In turn, behavioral and age differences have been associated with spatial functional topography rather than dynamic differences within individual RSNs (Bijsterbosch et al., 2018; Bozek et al., 2018; Kong et al., 2019), meaning that improved alignment between subjects is crucial to improve the accuracy of behavior or age models.

### 4.3 Methodological considerations

The selection of the number of independent components in the group ICA was based on previous work with this cohort (Eyre et al., 2021; Molloy and Saygin, 2022; Nielsen et al., 2022) as well as adult parcellations (Gordon et al., 2016; Yeo et al., 2011). While this collection of studies includes both hard and soft parcellations ranging from 7 to 30 components, the exact number that strikes a balance between functional homogeneity and interpretability remains an open debate (Gordon et al., 2016). For consistency with the latest studies on this dataset, where 30 components were used to describe volumetric RSNs (Eyre et al., 2021) that include subcortical structures, we opted for 20 components for our surface-based analysis excluding subcortical regions. As a result, we find that both group ICA and the individual parcellations align with biologically relevant areas previously associated with increased neuronal activity in adult task-based experiments (Glasser et al., 2016; Nickerson, 2018; Smith et al., 2009).

The algorithm and imaging modality chosen to co-register individual cortical surfaces proved another crucial methodological question in this study. Inter-subject feature alignment is known to gravely affect functional comparisons of the brain, a problem that is only exacerbated during the dramatic growth seen in the perinatal period. Even when alignment is satisfactory, individual differences have been shown to obscure important functional areas in group-level averages (Gordon et al., 2022, 2017). When first started this study using MSM registration method optimized for folding alignment (Robinson et al., 2018), some clusters of higher order networks were not strongly present in group ICA but seemed to emerge in many individual ICs. We then optimized registration for functional alignment by increasing the penalization for surface distortion, a method named ‘MSMSulc’ in (Williams et al., 2023). Consequently, group ICA included a complete and strong default mode network, illustrating the relative sensitivity of these analyses to cortical alignment across subjects.

As a alternative strategy, a recent study on neonatal population has explored multimodal alignment methods that supplement cortical folding-driven alignment with functional connectivity gradients to refine inter-subject registration (Wang et al., 2023), echoing previous efforts in adults (Glasser et al., 2016). In the present study, however, the total number of scans was significantly lower and each individual is represented by shorter total fMRI timeseries, making us less confident in the sharpness of individual FC gradients to drive further alignment. Additionally, the multiband factor of 9 used in the dHCP acquisition results in a non-homogeneously lower SNR that complicate this method further (Risk et al., 2021). Being a non-trivial problem, we deemed multimodal registration of the individuals in this dataset to be the scope of future research.

Head movement modulated by arousal states may influence the resting-state functional connectivity patterns in infants (Mitra et al., 2017). Indeed, it is plausible that certain infants may exhibit increased alertness or be in a state of active sleep, leading to increased head movement compared to other infants who are in a quiet sleep state (Denisova, 2019). Applying an aggressive frame censoring approach that selectively retains BOLD data segments with minimal motion can introduce a potential systematic bias towards subjects or data segments associated with a particular state. Thus, to mitigate any potential bias influenced by the arousal state of the infants, we implemented a conservative frame censoring approach. This approach guarantees an equal number of consecutive frames for each subject, ensuring that the segments remain qualitatively and quantitatively comparable across the entire cohort.

## 5 Conclusions

In this study, we employed a surface-based hierarchical Bayesian framework to leverage shared information across subjects and produced cleaner estimates of individual functional connectivity maps in neonates when compared to classic RSFC analyses. Improved detection of individual differences was evidenced by a maturational path in almost all brain regions. Importantly, our work extends beyond existing research by showing that significant maturational changes are not only restricted to primary sensory networks but also present in higher-order networks in the first weeks of postnatal life. These results illustrate the potential advantages of combining surface-based processing and template-based approaches to inform individual variability in very young populations, opening the door to precision neuroimaging studies of early brain development with enhanced accuracy and reliability.

## 6 Data and code availability

The neonatal data in this study is part of the second release of the developing Human Connectome Project and are available to download (https://www.developingconnectome.org). The cortical atlas is also publicly available (https://brain-development.org/brain-atlases/atlases-from-the-dhcp-project/cortical-surface-atlas-bozek/) . The in-house code used in this study includes shell scripts, python scripts, and R implementations of the templateICAr (https://github.com/mandymejia/templateICAr) and ciftiTools (Pham et al., 2022) libraries. A version of the code, edited for clarity, is available and maintained at https://github.com/FerradalLab/babyBayes.

## 7 Author contributions

S.L.F. developed the original idea and research question. S.L.F. and D.D. designed the methodology. A.F.M. and D.D.P. developed the original methods and tools. D.D. designed, coded, and ran the analysis. D.D. and S.L.F. interpreted the results and wrote the manuscript. All authors contributed to editing the manuscript.

## 9 Acknowledgments

Data were provided by the developing Human Connectome Project, KCL-Imperial-Oxford Consortium funded by the European Research Council under the European Union Seventh Framework Programme (FP/2007-2013) / ERC Grant Agreement no. [319456]. We are grateful to the families who generously supported this work. The non-dHCP pipeline analyses were performed on the Indiana University HPC Cluster (https://uits.iu.edu/). Template ICA was developed via grant support from the National Institute of Biomedical Imaging and Bioengineering (R01 EB027119).

## Footnotes

1 dHCP Data Release 2019: https://drive.google.com/file/d/197g9afbg9uzBt04qYYAIhmTOvI3nXrhI/view

2 https://github.com/ecr05/dHCP_template_alignment.

