## Supplementary Material for "Individual patterns of functional connectivity in neonates as revealed by surface-based Bayesian modeling"

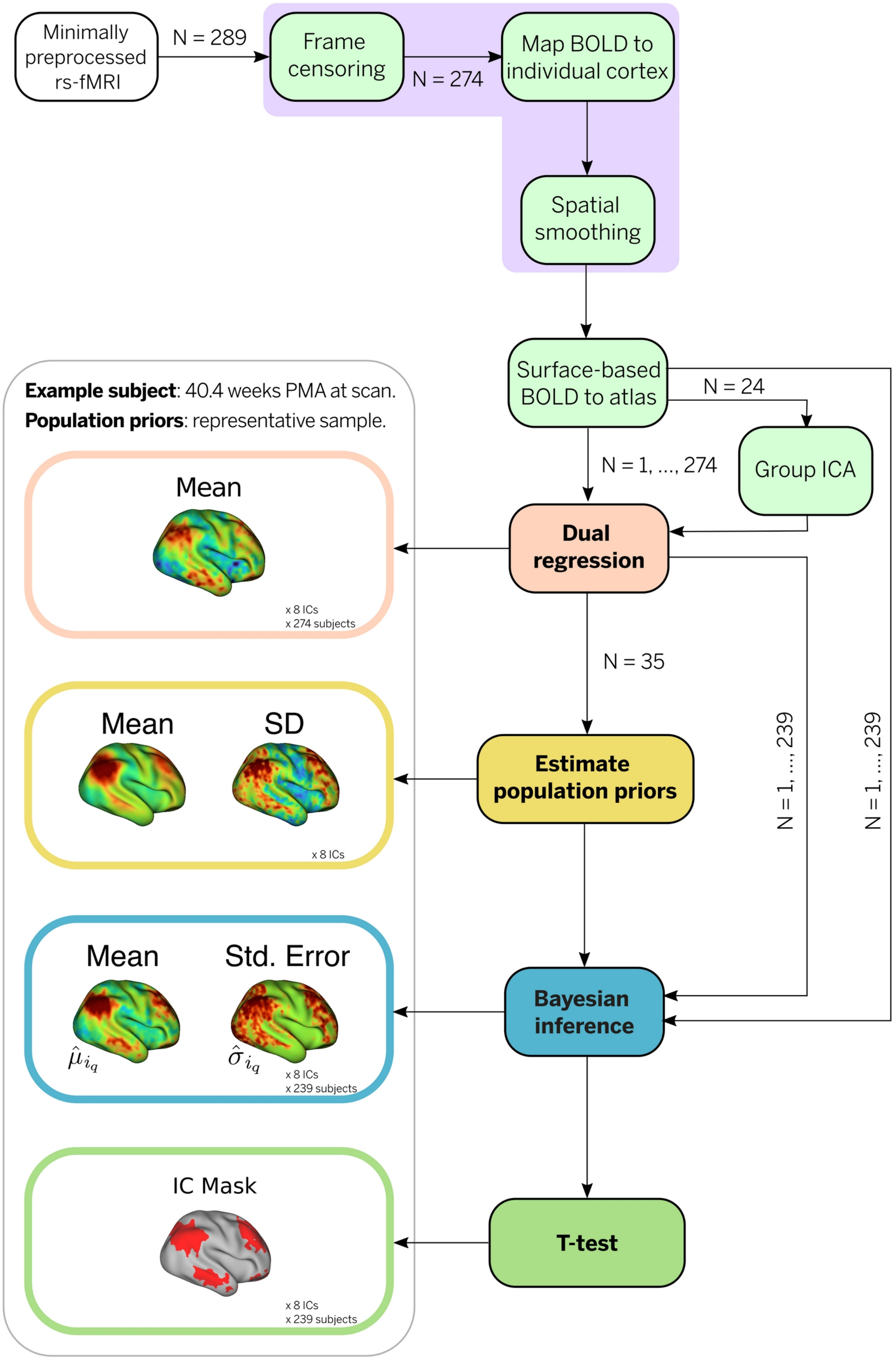


Figure S1: **Outline of the processing steps**. The minimally preprocessed BOLD volumes were projected onto their corresponding cortical surface and subsequently mapped to atlas space using spherical alignment. Group ICA maps were estimated from a subset of 24 neonates (age at scan: 43.5 - 45 weeks PMA). Rough estimates of individual IC maps were obtained for the whole cohort using dual regression. A population template (mean and inter-subject variance) was obtained from 35 subjects, uniformly sampled based on age at scan to provide an unbiased representation of the population. Using Bayesian inference, individual maps of mean and variance were estimated for all infants, excluding the 35 subjects considered in the estimation of the population priors. Individual t-statistic maps were estimated for each subject and significant areas of engagement were identified via t-test ($p < 0.05$).

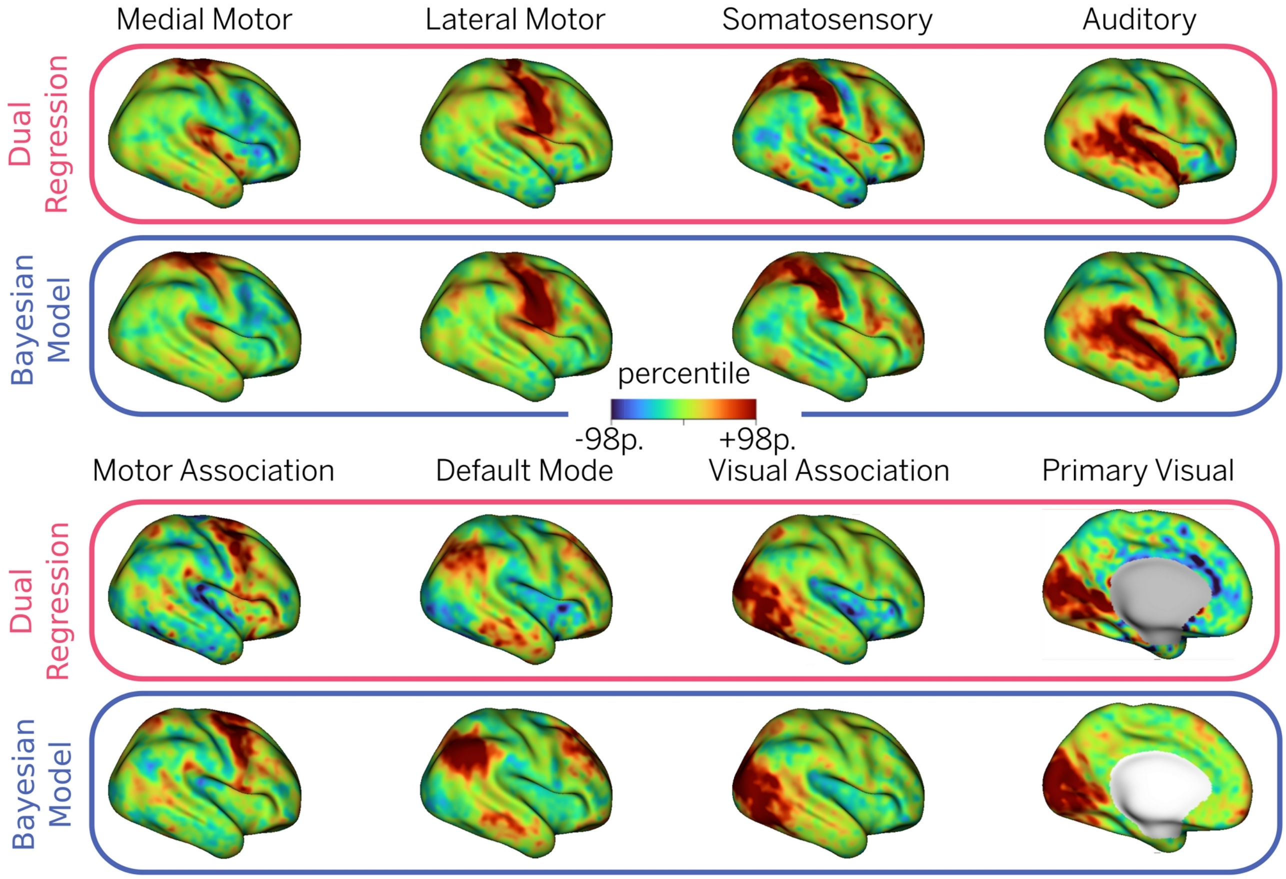


Figure S2: **Full comparison of cortical Bayesian IC maps and dual regression for a single subject**. Eight networks obtained for a term-born neonate (age at birth: 40 weeks gestational age) scanned at 42.6 weeks PMA. All networks were projected onto an inflated 40-week surface atlas.

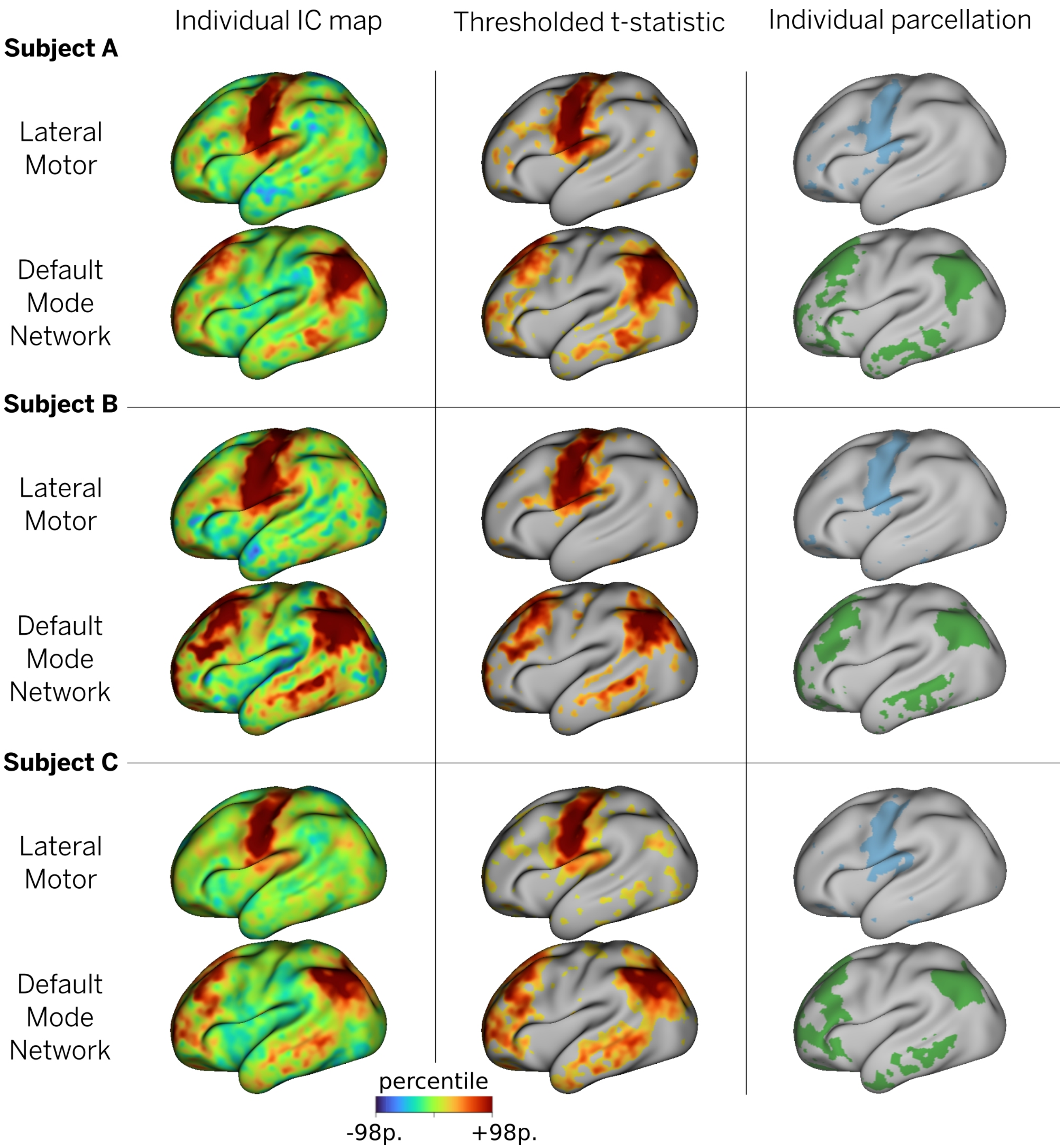


Figure S3: **Construction of individual parcellation**. For the three subjects in Fig. 3, a step-by-step example of the parecellation is given for two different components. Left column: the individual estimate of FC is obtained from Bayesian inference (see Fig. S1). Middle column: a t-test is performed and each network is thresholded at a statistically significant level ($p < 0.01$). Right column: each vertex is assigned to the highest t-statistic value after thresholding. The finalized parcellation is depicted in Fig. 3.

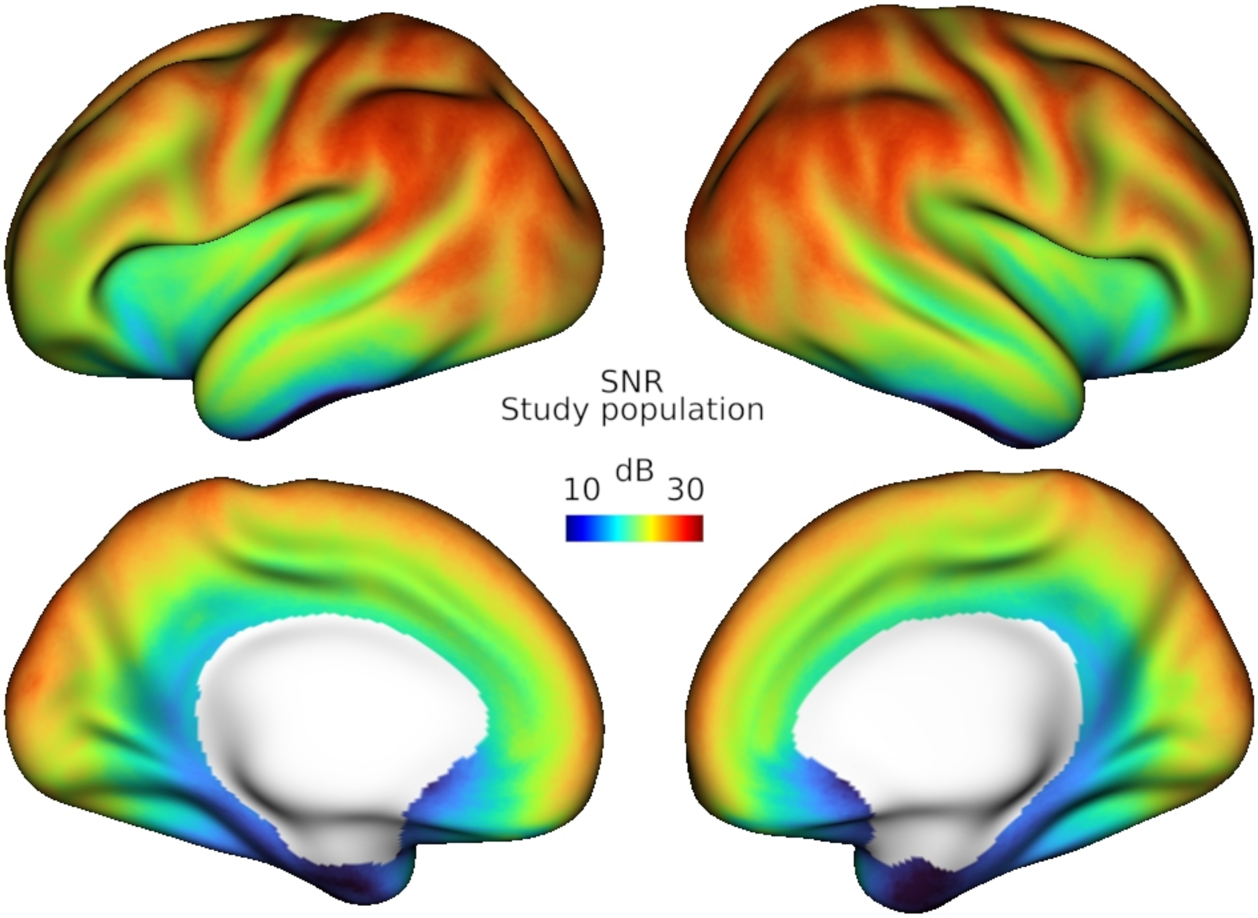


Figure S4: **Global SNR.** Signal-to-noise ratio (SNR) for the entire cohort (N = 274) computed as the ratio between the mean BOLD signal and the mean standard deviation at each vertex. SNR was calculated after data preprocessing (including frame censoring, volume-to-surface mapping and spatial smoothing) and projected onto the inflated 40-week surface atlas.

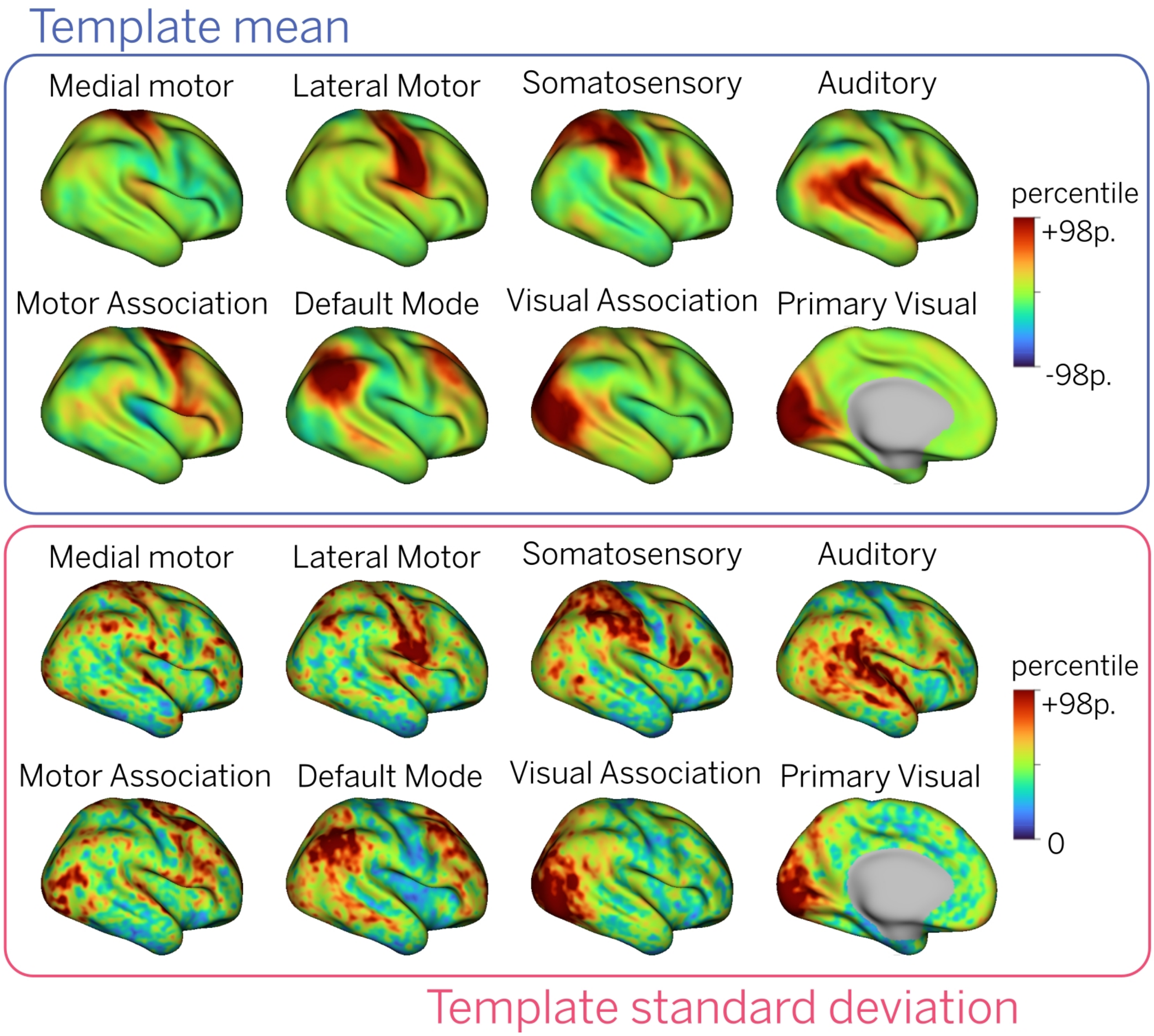


Figure S5: **Empirical population priors or template for fourteen RSNs**. Mean and standard deviation maps obtained from a representative subset of term-born infants from the dHCP database. All maps were projected onto an inflated 40-week surface atlas.

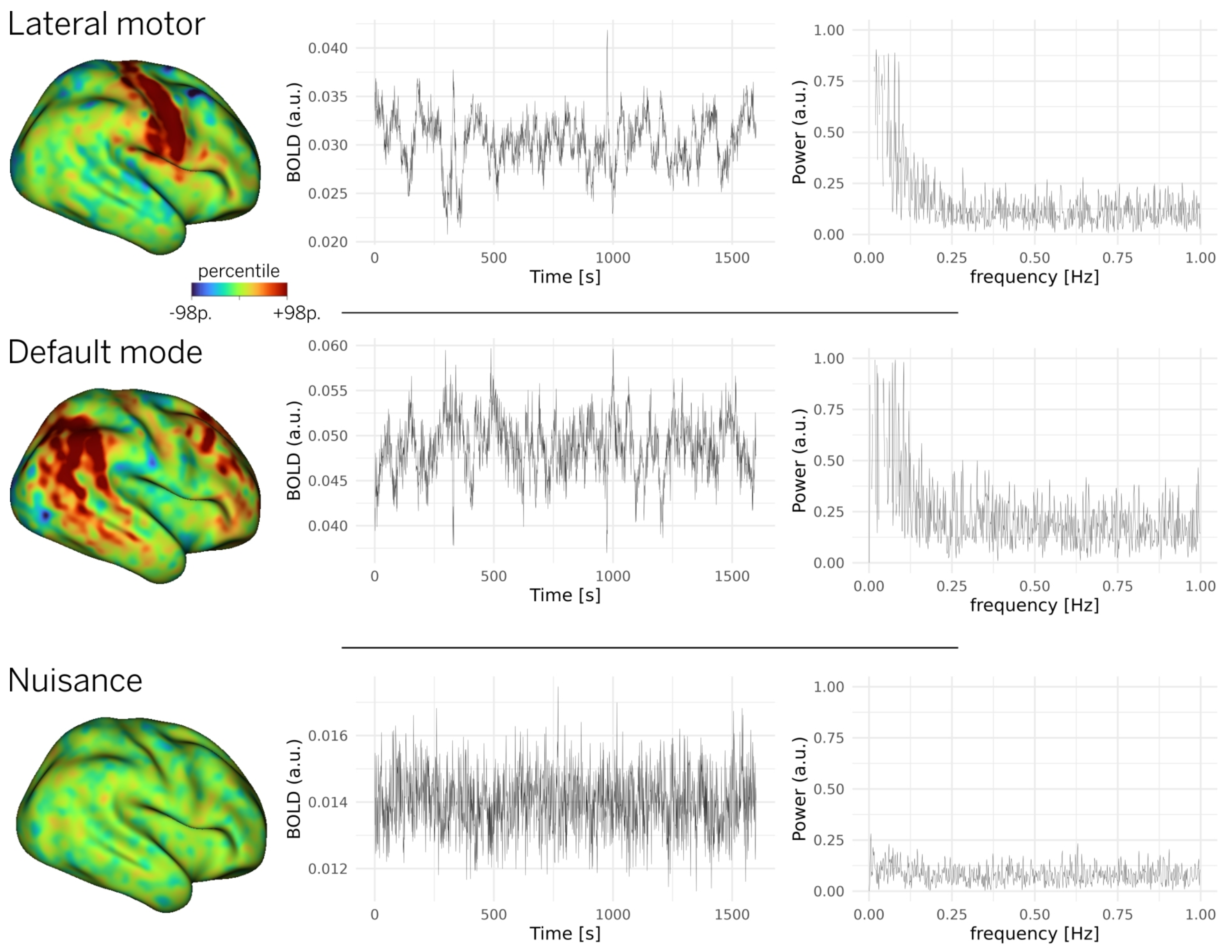


Figure S6: **Example of signal and nuisance group ICs**. Regressed IC maps, timeseries, and power spectra for a single subject. Left column: regressed IC map. Middle column: regressed timeseries of the IC, also known as column of the mixing matrix. Right column: power spectrum associated with the timeseries.

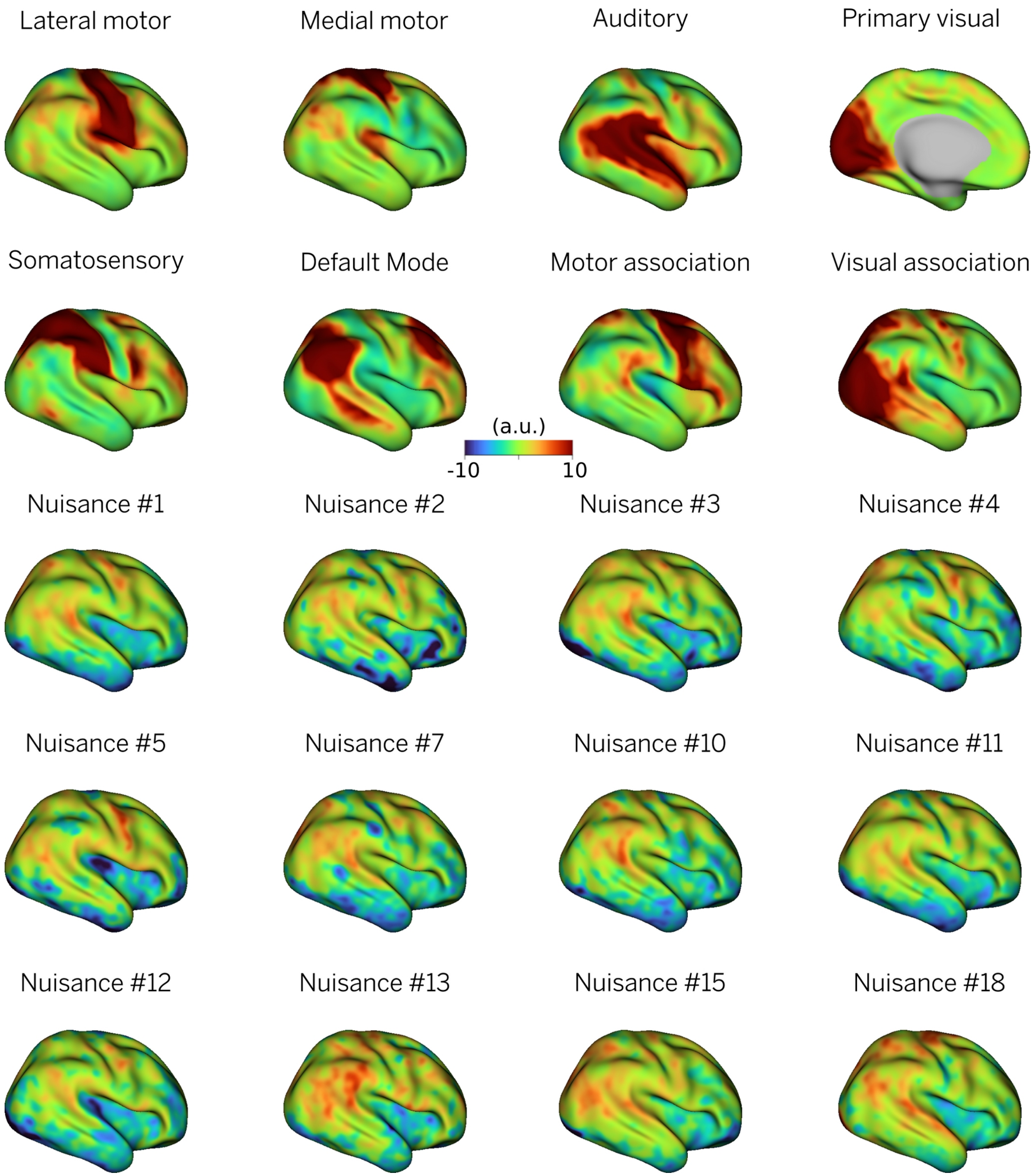


Figure S7: **Results of group ICA**. All the 20 independent components obtained from group ICA for a subset of 24 subjects of the dHCP dataset are depicted on the 40-week inflated atlas. The first eight components were designated as “signal” and used for the subsequent analyses. The rest were deemed “nuisance” and were not used.
